# Identification of functional immune and neuronal tumour cells in glioma

**DOI:** 10.1101/2022.11.05.515316

**Authors:** Rachel Naomi Curry, Malcolm F. McDonald, Qianqian Ma, Jochen Meyer, Isamu Aiba, Brittney Lozzi, Alexis Cervantes, Yeunjung Ko, Estefania Luna-Figueroa, Dong-Joo Choi, Zhung-Fu Lee, Junzhan Jing, Arif O. Harmanci, Anna Rosenbaum, Peihao He, Carrie Mohila, Ali Jalali, Jeffrey Noebels, Xiaolong Jiang, Benjamin Deneen, Ganesh Rao, Akdes Serin Harmanci

**Affiliations:** Graduate School of Biomedical Sciences, Baylor College of Medicine, Houston, TX; Center for Cell and Gene Therapy, Baylor College of Medicine, Houston, TX; Medical Scientist Training Program, Baylor College of Medicine, Houston, TX; Development, Disease Models, and Therapeutics, Baylor College of Medicine, Houston, TX; Center for Cancer Neuroscience, Baylor College of Medicine, Houston, TX; Department of Neurology, Baylor College of Medicine, Houston, TX; School of Biomedical Informatics, University of Texas Health Science Center, Houston, TX; Department of Pathology, Texas Children’s Hospital, Houston, TX; Department of Neurosurgery, Baylor College of Medicine, Houston, TX; Jan and Dan Duncan Neurological Research Institute, Texas Children’s Hospital, Houston, TX; Department of Human and Molecular Genetics, Baylor College of Medicine, Houston, TX

## Abstract

Despite advances in molecular profiling, therapeutic development has been hindered by the inability to identify and target tumour-specific mechanisms without consequence to healthy tissue. Correspondingly, a computational framework capable of accurately distinguishing tumour from non-tumour cells has yet to be developed and cell annotation algorithms are unable to assign integrated genomic and transcriptional profiles to single cells on a cell-by-cell basis. To address these barriers, we developed the Single Cell Rule Association Mining (SCRAM) tool that integrates RNA-inferred genomic alterations with co-occurring cell type signatures for individual cells. Applying SCRAM to glioma, we identified tumour cell trajectories recapitulate temporally-restricted developmental paradigms and feature unique co-occurring identities. Specifically, we validated two previously unreported tumour cell populations with immune and neuronal signatures as hallmarks of human glioma subtypes. *In vivo* modeling revealed a rare immune-like tumour cell population resembling antigen presenting cells can direct CD8+ T cell responses. In parallel, Patch sequencing studies in human tumours confirmed that neuronal-like glioma cells fire action potentials and represent 40% of *IDH1* mutant tumor cells. These studies identified new glioma cell types with functional properties similar to their non-tumour analogues and demonstrate the ability of SCRAM to identify these cell types in unprecedented detail.

## Introduction

In the era of cancer genomics, the advent of high-throughput single cell sequencing technologies has allowed for examination of cellular heterogeneity in genomic, transcriptomic and epigenomic detail^1^. Studies employing these technologies have demonstrated dynamic cellular and subcellular hierarchies that are spatiotemporally distinct and have revealed transcriptional profiles that can be utilized to classify tumours into clinically-relevant molecular subtypes^2–5^. Despite these advances, the ability of sequencing pipelines to reliably identify tumour cells and the co-occurring genomic and transcriptomic cell states that define them has yet to be attained. Within malignant glioma, the most lethal form of brain tumors, resolution of this complexity represents a major impediment to therapeutic development that is further compounded by a diverse cellular constituency, complex array of tumour-specific genetic variants and inability to distinguish glioma cells from their non-tumor counterparts. To this end, we have developed a novel single cell computational tool, Single Cell Rule Association Mining (SCRAM) capable of accurately identifying tumour cells and defining co-occurring cellular states for individual cells. SCRAM integrates three orthogonal features to identify tumour cells in single cell resolution: (1) cell type transcriptional profiling; (2) RNA-inferred copy number variant (CNV) calling; and (3) RNA-inferred single nucleotide variant (SNV) calling. Our studies revealed that more than half of tumour cells feature transcriptional profiles matching more than one cell type, including an infrequent astrocytoma-specific population with transcriptional and functional properties of antigen presenting immune cells. In parallel, SCRAM uncovered that the majority of *IDH1* mutant (IDH1^mut^) tumour cells possess GABAergic neuronal-like transcriptomes and fire action potentials. Our observations demonstrate that tumour cells endowed with immune and neuronal expression profiles can acquire functional cellular states and highlight the utility of using our computational framework for characterizing pseudo cell types in cancer.

## Results

### SCRAM reliably discriminates between tumour and non-tumour cell types

To validate SCRAM as a reliable computational framework (**Fig. 1A**), we examined whether SCRAM annotation was consistent with established transcriptional profiles for previously published human and mouse single-cell RNA-sequencing (scRNA-seq) datasets. We manually curated a cell type marker list for assigning cell identities based on at least two cell type–specific markers reported by EnrichR^6^, Allen Brain Atlas^7,8^, and other studies (**Fig. S1, Tables S1**,**2**). SCRAM could assign cell type identities to greater than 75% of cells across eight scRNA-seq datasets (**Fig. 1B**) and identified greater than 83% of cells as possessing transcriptional profiles meeting the criteria for multiple cell types (**Fig. 1C**). Using established glioma transcriptional profiles^5,9^ and our manually curated marker list, we assigned cell type annotation to individual cells and found that SCRAM-assigned identities were consistent with those defined in previous reports (**Figs. 1D–G**). Our analysis found that greater than two-thirds of sequenced cells exhibited co-occurring cell type annotations, suggesting that glioma cells possess transcriptional signatures consistent with multiple cellular states. Further validation using the 10X Genomics spatial transcriptomics dataset of *IDH1–*wild-type (IDH1^WT^) glioma revealed that various permutations of co-occurring cellular states exist within single-cell clusters, demonstrating that clusters encompass a heterogenous group of cell types with various genomic and transcriptomic profiles (**Figs. 1H,I; S2A**). Because SCRAM can define both transcriptional signatures and RNA-inferred genomic CNVs, we were able to further characterize the respective lineages of tumour cells. Our results confirmed that tumour cells are spatially dispersed throughout the non-tumour landscape and present with diverse transcriptomic cellular profiles within a small anatomical region. These analyses validated the ability of SCRAM to replicate previously defined cellular annotations and demonstrated the ability of SCRAM to resolve intra-tumoural heterogeneity by defining co-occurring cellular identities and cellular states with high resolution.

**Figure 1.**
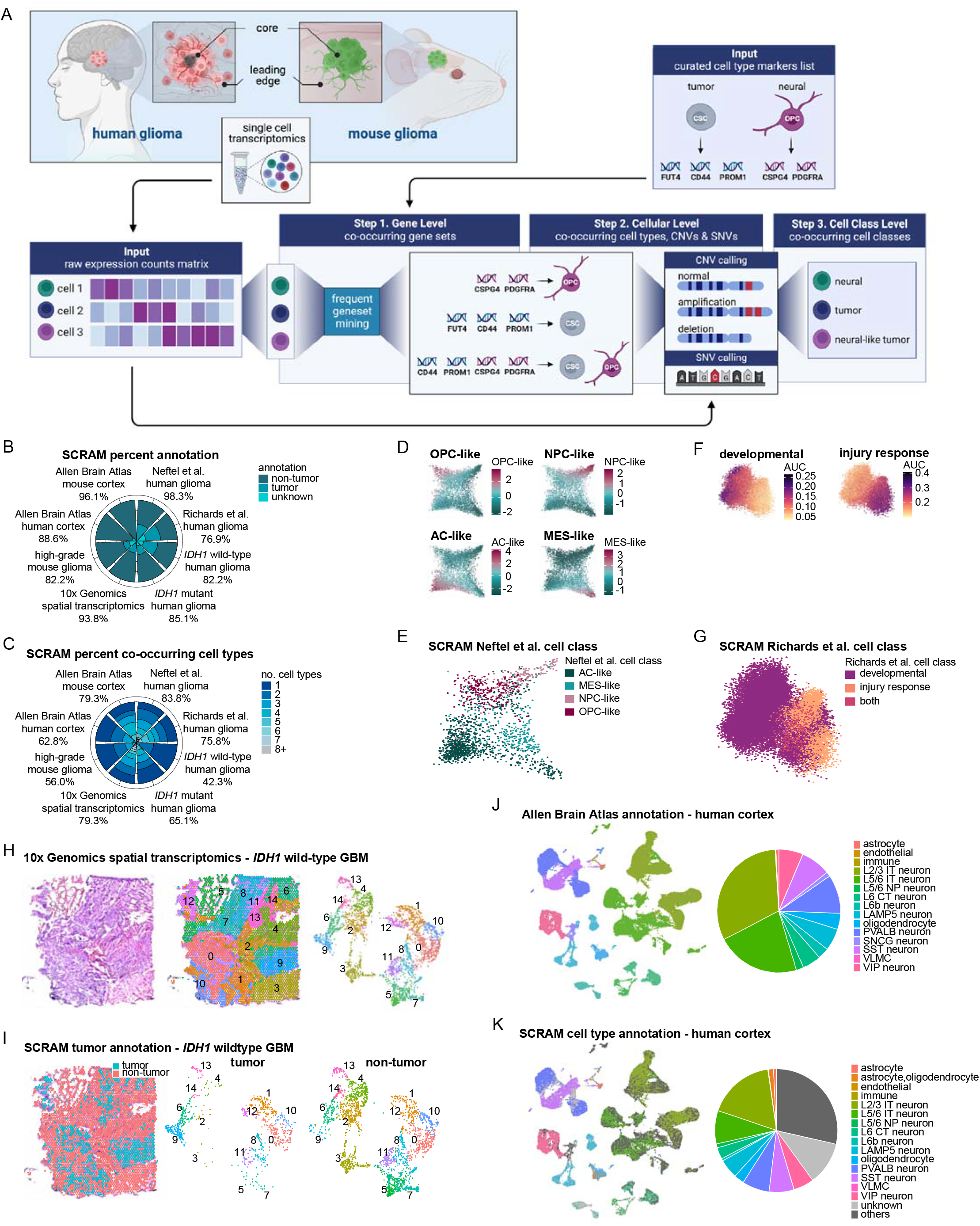
SCRAM reliably assigns cell type annotations to single-cell RNA-sequencing datasets on a cell-by-cell basis. (A) Schematic representation of the Single-Cell Rule Association Mining (SCRAM) pipeline. (B) Percentages of SCRAM-annotated cells assigned to non-tumour, tumour, or unknown cell types across eight datasets. Percentages shown below dataset names indicate the total cell percentages assigned to each cell type annotation using SCRAM. (C) Percentages of SCRAM-annotated cells by the number of co-occurring cell types. Percentages shown below dataset names indicate cells with more than one cell type annotation. (D) Reproduction of meta-modules from Neftel et al.^5^, showing astrocyte-like (AC-like), oligodendrocyte precursor cell–like (OPC-like), mesenchymal-like (MES-like), or neural precursor cell–like (NPC-like) cellular states. (E) Meta-module of SCRAM-identified tumour cells from Neftel et al. dataset. (F) Reproduction of cell state modules from Richards et al.^9^, showing developmental and injury response programmes. (G) SCRAM-assigned developmental and injury response cell classes for the Richards et al. dataset. (H) Haematoxylin and eosin staining and corresponding Seurat clusters are shown for the 10X Genomics spatial transcriptomics *IDH1–*wild-type (IDH1^WT^) glioblastoma (GBM) dataset. (I) SCRAM-assigned tumour and non-tumour annotations for the 10X Genomics spatial transcriptomics IDH1^WT^ GBM dataset show tumour cells are embedded throughout non-tumour tissue. Seurat clusters separated by tumour and non-tumour annotations show clusters comprised of tumour and non-tumour cells. (J) Cell type annotation for human cortex dataset provided by Allen Brain Atlas. (K) SCRAM-assigned cell type annotation for the Allen Brain Atlas human cortex dataset reveals co-occurring cell identities (dark grey) and unknown transcriptional cell states (light grey).

After confirming the capacity of SCRAM to accurately assign cellular identities, we assessed the ability of SCRAM to distinguish tumour cells from non-tumour cells. We analysed non-tumour human and mouse cortex scRNA-seq datasets from the Allen Brain Atlas and found that SCRAM reliably distinguished tumour from non-tumour cells with greater than 99% accuracy across species. SCRAM was able to annotate greater than 88% of cells in these datasets, replicating 60% and 86% of the Allen Brain Atlas annotations for the human and mouse datasets, respectively (**Figs. 1J,K; S2B,C**). Individual cellular annotations revealed that clusters comprised heterogeneous cell types, and we further refined these expression signatures using the *de novo* marker function in SCRAM (**Figs. S2D–G**). These data demonstrate that SCRAM reliably distinguishes between tumour and non-tumour cell types and accurately assigns cell identities in humans and mice. Furthermore, these studies highlighted the utility of SCRAM for examining single-cell expression profiles and characterizing cellular heterogeneity in the mammalian brain.

### SCRAM elucidates novel developmental tumour cell states in glioma

Having validated SCRAM, we next aimed to define the genomic and transcriptomic landscape of tumour cells in our integrated scRNA-seq dataset consisting of 195,063 cells from seven malignant glioma patients (**Fig. S3A; Table S3**). SCRAM analyses employing the CaSpER^10^ CNV-calling algorithm (**Figs. S3B**,**C**) confirmed that IDH1^WT^ tumours had increased CNV incidence (**Figs. 2A–D; S3D**,**E**), whereas chromosome 1p19q co-deletions were the predominant alterations in IDH1^mut^ oligodendroglioma. Parallel analyses using XCVATR^11^ to call SNVs validated that IDH1^mut^ tumours featured *IDH1* mutations, whereas IDH1^WT^ tumours featured *EGFR, TP53*, and *PTEN* mutations (**Figs. 2E**,**F; S3F**,**G**). Although DNA-based sequencing studies have described a higher mutational burden in IDH1^WT^ tumours, our RNA-inferred analyses identified more frequent SNV transcription at the mRNA level in IDH1^mut^ glioma than in IDH1^WT^ glioma. Of particular interest was the high incidence of mutations in a transcript mapping to the polycomb repressive complex 2 (PRC2) *SUZ12* sequence. Further examination revealed that the mutated *SUZ12* sequence is identical to the *SUZ12* pseudogene (*SUZ1*2*P1*) sequence. This transcript was detected at high levels in all glioma patients, suggesting either that *SUZ12* mutations are the most transcribed SNVs in our cohort or that *SUZ12P1* is actively transcribed in glioma patients (**Figs. 2G,H; S3H**). Prior reports have observed hypermethylation of PRC2 regions in malignant tumours, which is pronounced in IDH1^mut^ subtypes^12,13^ and has identified mutant *SUZ12* as a driver of some peripheral nervous system malignancies^14,15^. Whether high transcriptional penetrance of *SUZ12P1* or mutated *SUZ12* mechanistically contributes to malignant glioma warrants future mechanistic inquiries.

**Figure 2.**
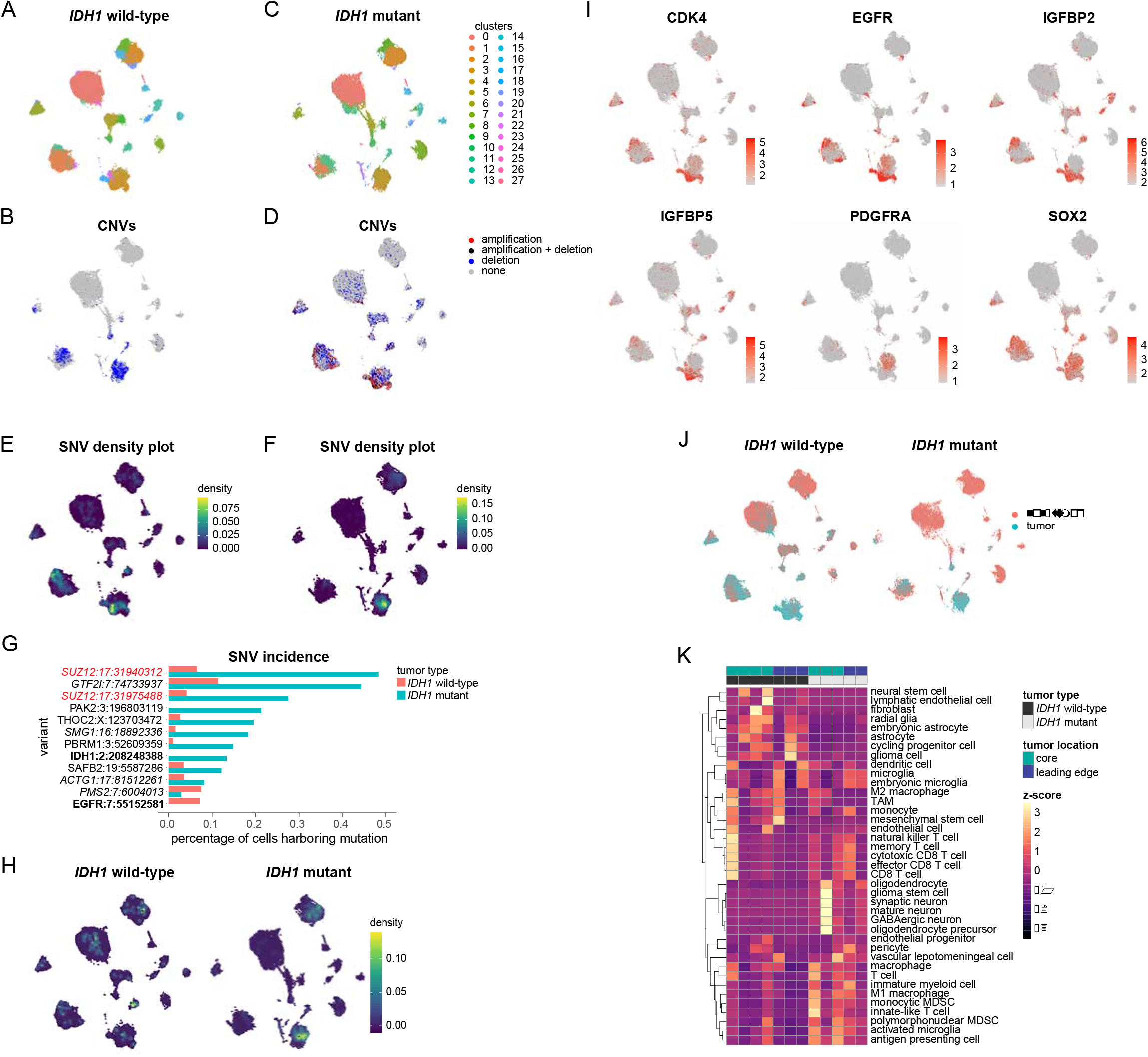
Integrated RNA-inferred genomic and transcriptomic cellular states reveal unique molecular signatures in *IDH1–*wild-type and *IDH1*-mutant glioma. (A) Seurat clusters are shown for 157,316 cells from *IDH1* wild-type (IDH1^WT^) glioma patients (n=4). (B) Feature plot showing total copy-number variant (CNV) calls by Single-Cell Rule Association Mining (SCRAM) for IDH1^WT^ tumours. (C) Seurat clusters are shown for 37,747 cells from *IDH1-* mutant (*IDH1*^*mut*^) glioma patients (n=3). (D) Feature plot showing total CNV calls by SCRAM for IDH1^mut^ tumours. (E) Density plot showing single-nucleotide variant (SNV) density for rare and COSMIC variants in IDH1^WT^ glioma. (F) Density plot showing SNV density for rare and COSMIC variants in IDH1^mut^ glioma. (G) Bar plot of the top 12 most frequent SNVs. Plot shows the percentage of cells harbouring rare and COSMIC SNVs in IDH1^WT^ and IDH1^mut^ glioma; italics denote genes with known pseudogenes; bold denotes known driver genes in glioma. (H) Density plot showing either *SUZ12* mutations or the *SUZ12P1* pseudogene detected in IDH1^WT^ and IDH1^mut^ samples. (I) Feature plots for tumour cell markers used to assign tumour annotation. (J) SCRAM-assigned tumour annotations are shown for IDH1^WT^ and IDH1^mut^ glioma. (K) Heatmap showing enrichment for SCRAM-assigned cell types by tumour type and sample location; z-score for expression of cell types is shown.

Using CNV and SNV annotation, together with established tumour marker expression (**Fig. 2I; Figs. S4A–F**), cells were assigned tumour or non-tumour status (**Fig. 2J**). Tumour cells in both IDH1^WT^ and IDH1^mut^ samples primarily mapped to one of three SCRAM-assigned cell classes (embryonic-neural, embryonic, or neural) and one of four co-occurring cell lineages (glial, glial-neuronal, myeloid, or neurodevelopmental), emphasizing shared transcriptomic features among glioma subtypes (**Figs. S5A–D**). An examination of cell types by patient revealed within- and between-patient heterogeneity and illuminated various myeloid-derived immune cell signatures within IDH1^mut^ tumour core samples (**Fig. 2K; S5A–D**). Importantly, CNVs were the only marker in 17% of tumour cells, representing a tumour cell population that could not be assigned to any cell type (**Fig. S5E**). Using the SCRAM *de novo* marker function, we identified markers of these unclassified tumour cells, which include *TRIO, RNF180* and *ELL2* for cells with chromosome 5 and 7 gains, and *STAU1, NAA20* and *DSTN* for cells with chromosome 20 gains (**Table S4**). Since *de novo* marker genes are differentially expressed in CNV-bearing tumor cells, they may serve as putative tumor-specific markers of glioma cells and should be further studied in future scientific endeavours. Collectively, these data demonstrate that IDH1^WT^ and IDH1^mut^ tumours are defined by unique, co-occurring genomic and transcriptomic profiles that exist in varying proportions among glioma subtypes and highlight the utility of SCRAM for defining transcribed mutations and integrated cellular states in human glioma.

Previous observations have indicated that spatially-distinct regions within individual tumours possess unique transcriptomic and genomic signatures^16^; therefore, we sought to spatially resolve the co-occurring cell types identified in our glioma dataset by comparing tumor cells surgically resected from the leading edge with those of the tumor core. Analysis of each Seurat-generated cluster revealed that clusters were diverse and frequently featured both tumour and non-tumour cells with similar transcriptional expression patterns (e.g., astrocytes and astrocyte-like tumour cells). The largest tumour cell cluster in our human dataset (Cluster 1) contained cells that were uniformly annotated as astrocytes and featured various co-occurring chromosome 5p, 5q, 7p, 7q, and 17q amplifications and chromosome 10p, 10q, and 22q losses (**Figs. 3A–D**). Visualization of co-occurring CNVs by patient showed that IDH1^WT^ tumour cells in the leading edge featured fewer variations of co-occurring chromosomal rearrangements than tumour cells in the core (**Fig. 3E**). This finding suggests that clonal tumour cells in the leading edge emerge from specific glioma cell subclones residing in the tumour core and is further supported by lineage-tracing studies showing that leading edge cells emanate from more primitive cell types in the tumour core (**Figs. 3F–H**). Using RNA velocity pseudotime analysis^17^, we found that these astrocytic tumour cells emanate from stem cell–like tumour cells in Cluster 23 and transition through an intermediate cell type found primarily in Cluster 3 **(Figs. S6A–D**). These intermediate cell types possess neurodevelopmentally restricted transcriptional profiles resembling those of embryonic astrocytes or radial glia while retaining the frequently co-occurring CNV annotations observed in Cluster 1. By contrast, the progenitor– and stem cell– like cells found in Cluster 23 feature only a single CNV occurrence and are characterized by the addition of primitive, co-occurring cell identities, including neural stem cell (NSC) and glioma stem cell (GSC), as well as oligodendrocyte annotation (**Figs. 3I–L**). These observations suggest that astrocyte-like tumour cells with widespread chromosomal anomalies derive from a chromosomally intact stem cell–like cell population featuring transcriptional profiles resembling those of oligodendrocytes. These cells transition through an intermediate cell state in which stem cell signatures are lost and pervasive co-occurring CNVs are gained (**Fig. 3M**). Synchronously, SCRAM identified analogous populations of glioma cells in our scRNA-seq dataset from our piggyBac transposase–based *in utero* electroporation (pB-IUE) glioma model, which yields green fluorescent protein (GFP)-labelled *de novo* tumours in immunocompetent mice (**Figs. S6E–H**). These combined analyses identified a compendium of individual cellular states identified by SCRAM that parallel neurodevelopmental trajectories and defined co-occurring spatiotemporal hierarchies.

**Figure 3.**
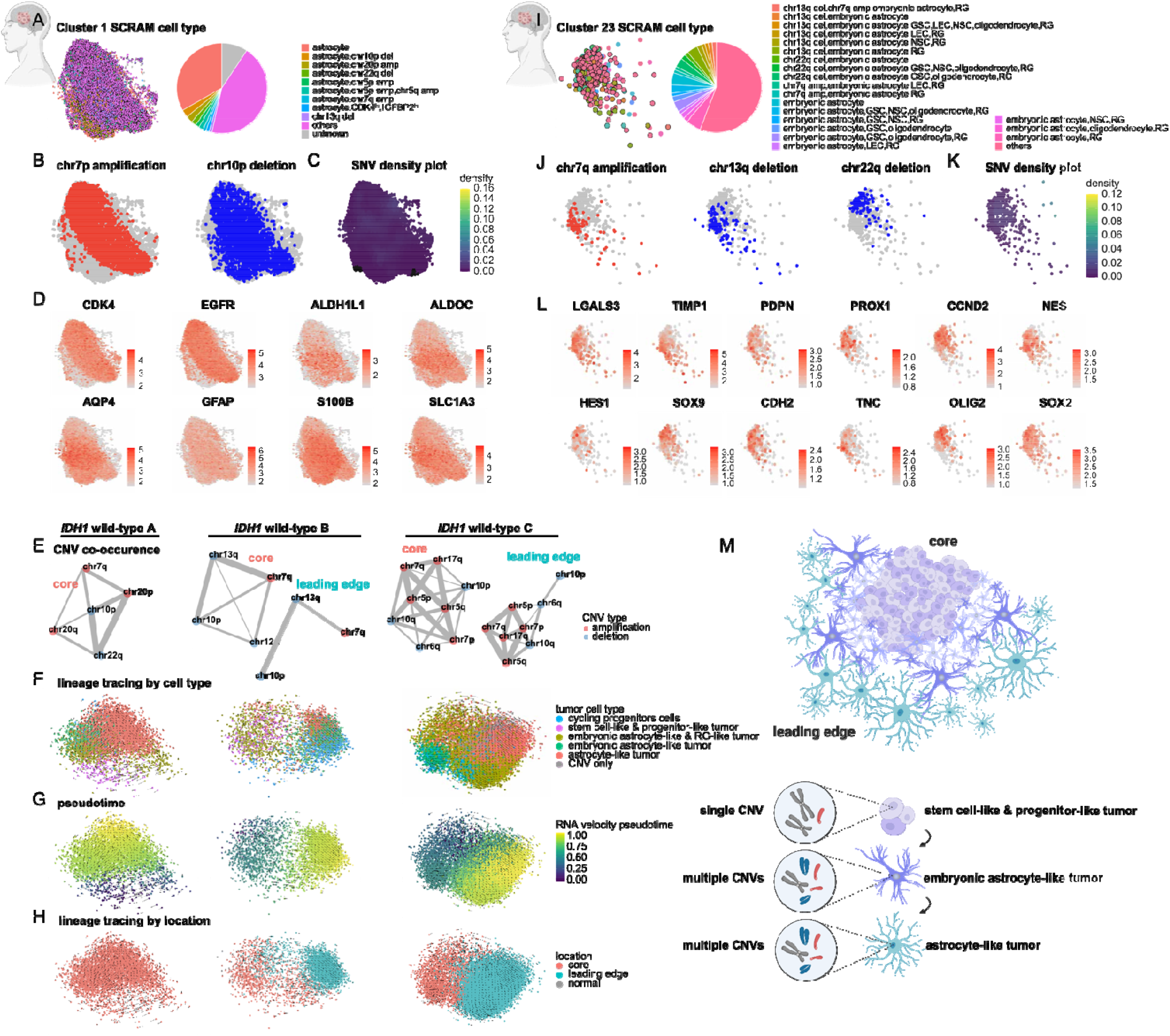
SCRAM uncovers new tumour cell states and developmental hierarchies in glioma. (A) Dimplot and corresponding pie chart showing Single-Cell Rule Association Mining (SCRAM)-identified astrocyte-like tumour cells for Cluster 1 in human glioma samples. Copy-number variant (CNV)-harbouring cells are outlined in black. (B) Feature plots showing exemplary large-scale CNVs. (C) Density plots showing single-nucleotide variant (SNV) density. (D) Feature plots for select tumour cell (*CDK4, EGFR*) and astrocyte (*ALDH1L1, ALDOC, AQP4, GFAP, S100B, SLC1A3*) markers. (E) Co-occurring CNVs for five patient-matched *IDH1*–wild-type (IDH1^WT^) glioma samples show leading edge cells emanate from specific co-occurring CNV lineages. Increased line thickness (weight) indicates more frequent co-occurrence. (F) RNA velocity lineage tracing by SCRAM-assigned cell type shows astrocyte-like tumour cells derive from stem-like and progenitor-like tumour cells and transition through an intermediate cell state marked by embryonic astrocytic and radial glial signatures. (G) RNA velocity pseudotime plots. (H) RNA velocity lineage tracing by tumour sample location shows leading edge cells are derived from glioma cells in the tumour core. (I) Dimplot and corresponding pie chart showing SCRAM-identified stem-like and progenitor-like cell type annotations for Cluster 23 cells in human glioma samples. CNV-harbouring cells are outlined in black. (J) Feature plots showing exemplary large-scale CNVs. (K) Density plot showing SNV density. (L) Feature plots for select embryonic astrocyte (*LGALS3, TIMP1*), GSC (*CCND2, OLIG2, SOX2*), LEC (*PDPN, PROX1*), NSC (*HES1, NES, SOX2, SOX9*), RG *(CDH2, HES1, NES, SOX2, TNC*) and oligodendrocyte (*OLIG2*) markers. (M) Schematic showing astrocyte-like tumour cells with multiple co-occurring CNVs in the leading edge emerge from embryonic astrocyte-like tumour cells with similar CNV profiles. Embryonic astrocyte-like tumour cells derive from stem-like and progenitor-like tumour cells in the core and feature single CNVs. GSC: glioma stem cell; LEC: lymphatic endothelial cell; NSC: neural stem cell; RG: radial glia.

### SCRAM reveals APC-like tumour cells in astrocytoma

Our prior analyses identified co-occurring neurodevelopmental cellular states within glioma cell transcriptomes, leading us to investigate whether these tumours also contain cellular signatures derived from non-neural lineages. We found 229 cells in Cluster 19 characterized by both tumour cell and APC annotations, which we termed APC-like tumour cells (ALTs). ALTs are rare (<1%) and unique to astrocytoma tumours in both IDH1^WT^ and IDH1^mut^ patients and frequently feature CNVs, cycling progenitor cell annotation, and high expression of the activated APC marker *CD83*^18^ and major histocompatibility complex (MHC) I and II genes^19^ (**Figs. 4A–D; Fig. S7A**). Many immune cell subtypes actively surveil the tumour microenvironment, where they engulf and display tumour antigens through processes of antigen presentation^20^. To confirm that ALTs were not artefacts representing immune cells phagocytosing tumour cells, we examined expression of the immune cell lineage marker *PTPRC* (CD45, **Fig. S7B**) and found no *PTPRC* expression in 69% of ALTs. SCRAM also identified an analogous population of *Ptprc-*negative ALTs in mouse glioma, which featured high *GFP* and *Cd83* expression (**Figs. S7C–E**). These data suggest the existence of a rare CNV-altered tumour cell population in high-grade astrocytoma, characterized by APC transcriptional signatures, and confirm the existence of a parallel cell population in our pB-IUE model.

**Figure 4.**
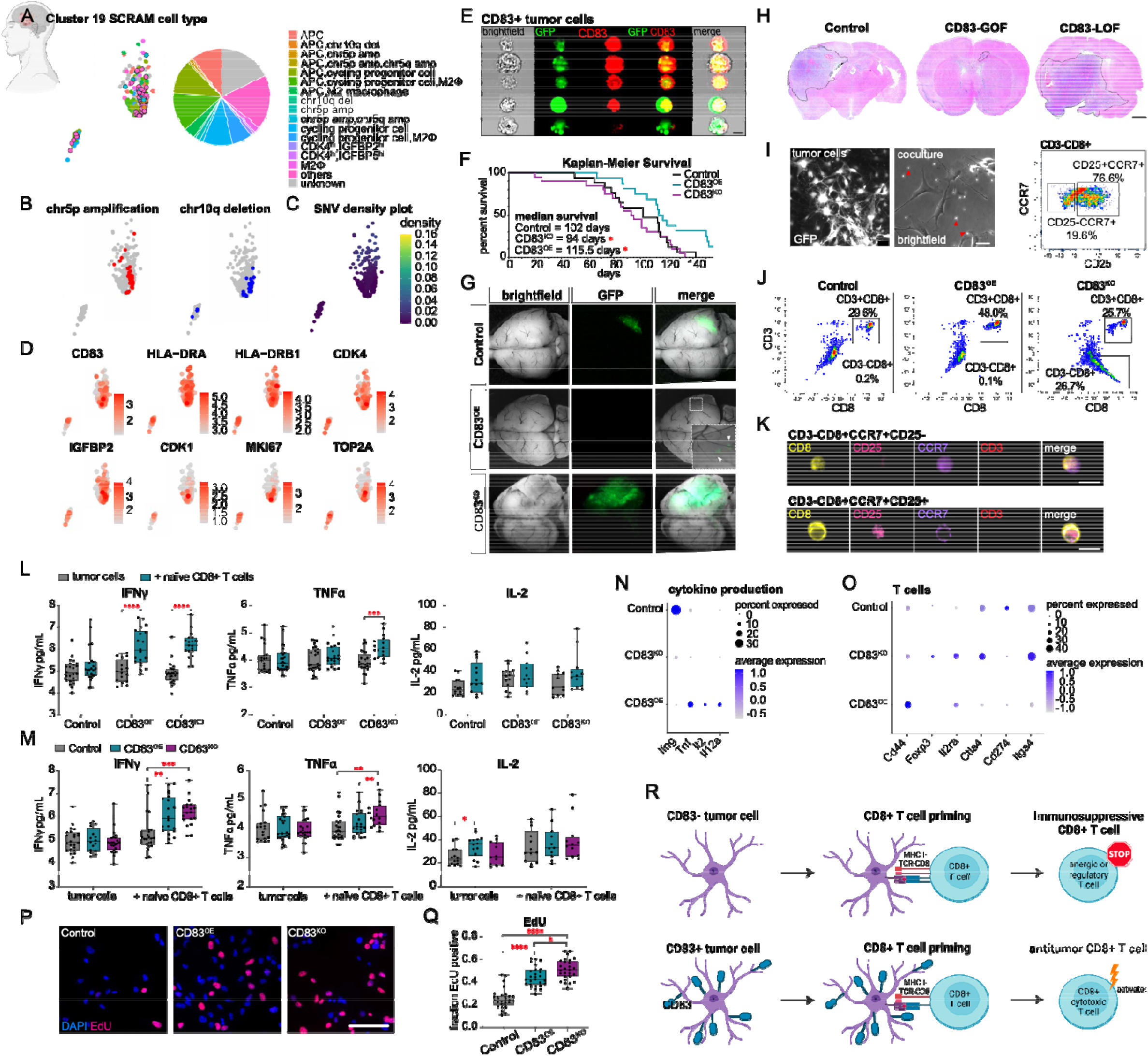
SCRAM-identified APC-like tumour cells mediate disease progression by restricting the immunosuppressive landscape in glioma. (A) Dimplot and corresponding pie chart showing Single-Cell Rule Association Mining (SCRAM)-identified antigen-presenting cell (APC)-like tumour cells (ALTs) for Cluster 19 in human glioma samples. Copy-number variant (CNV)-harbouring cells are outlined in black. (B) Feature plots showing exemplary large-scale CNVs. (C) Density plots showing single-nucleotide variant (SNV) density. (D) Feature plots for select tumour cell (*CDK4, IGFBP2*), cycling progenitor cell (*CDK1, CENPF, MKI67, TOP2A*), and APC (*CD83, HLA-DRA, HLA-DRB1*) markers. (E) Representative images from imaging flow cytometry of CD83^+^GFP^+^ ALTs demonstrating diffuse GFP expression throughout the cell; scale bar: 7 µm. (F) Kaplan–Meier survival analysis of mice bearing control (n=17), CD83 overexpressing (CD83^OE^; n=16), and CD83 knockout (CD83^KO^; n=20) tumours; \**p*<0.05. (G) Exemplary brightfield and GFP images from age-matched control, CD83^OE^, and CD83^KO^ tumours. (H) Representative haematoxylin and eosin whole-brain coronal sections from age-matched control, CD83^OE^, and CD83^KO^ tumours; black dotted lines denote tumour regions, scale bar: 1 mm. (I) Representative images from cell lines established from age-matched control, CD83^OE^, and CD83^KO^ tumours; scale bar: 20 µm. (J) Imaging flow cytometry plots showing an increased percentage of CD3^+^CD8^−^ T cells from CD83^OE^ (48.0%) co-cultures compared with controls (29.6%) and CD83^KO^ (25.7%) co-cultures. CD3^−^CD8^+^ T cells detected in 26.7% of CD83^KO^ T cells compared with 0.2% in control and 0.1% in CD83^OE^ T cells. Gating on CD3^−^CD8^+^ T cells from CD83^KO^ co-cultures shows two T cell populations: CD3^−^CD8^+^CCR7^+^CD25^−^ T cells (19.6%) and CD3^−^CD8^+^CCR7^+^CD25^+^ T cells (76.6%). (K) Representative images of CD3^−^CD8^+^CCR7^+^CD25^−^ and CD3^−^CD8^+^CCR7^+^CD25^+^ T cells from co-culture imaging flow cytometry experiments; scale bar: 7 µm. (L) Box and whisker plots for enzyme-linked immunosorbent assays (ELISA) show increased interferon-gamma (IFNγ) production in CD83^OE^ (\*\*\*\**p*<0.0001) and CD83^KO^ (\*\*\*\**p*<0.0001) co-cultures containing naïve CD8^+^ T cells. Tumour necrosis factor-alpha (TNFα) production only increased in CD83^KO^ co-cultures (\*\*\**p*<0.001); interleukin (IL)-2 production was unchanged. (M) Box and whisker plots for ELISAs show IFNγ production is increased in CD83^OE^ (\*\**p*<0.01) and CD83^KO^ (\*\*\*\**p*<0.001) T cell co-cultures relative to control co-cultures (\*\*\**p*<0.001). TNFα production increased in CD83^KO^ T cell co-cultures compared with CD83^OE^ (\*\**p*<0.01) and control cl-cultures (\*\**p*<0.01). IL-2 production increased in CD83^OE^ cells cultured in the absence of T cells compared with controls (\**p*<0.05). (N) Dot plots from control, CD83^OE^, and CD83^KO^ tumour single-cell RNA-sequencing (scRNA-seq) datasets show *in vivo* T cells enriched for *Ifng* (IFNγ) expression in control tumours and *Tnf* (TNFα), *Il2* (IL-2), and *Il12a* (IL-12) in CD83^OE^ tumours. (O) Dot plots from control, CD83^OE^, and CD83^KO^ tumour scRNA-seq datasets show T cells enriched for effector and memory T cell markers (*Cd44, Il2ra*) in CD83^OE^ tumours and anergic and regulatory T cell markers (*Foxp3, Il2ra, Ctla4, Itga4*) in CD83^KO^ tumours. (P) Representative images from EdU proliferation assays on control, CD83^OE^, and CD83^KO^ tumour cell lines *in vitro*. (U) Quantification of EdU+ cells for CD83^OE^ (\*\*\*\**p*<0.0001) and CD83^KO^ (\*\*\*\**p*<0.0001) cells, which are more proliferative than control cells; CD83^KO^ cells are more proliferative than CD83^OE^ cells (\**p*<0.05). (P) Schematic showing CD83 expression in tumour cells enables CD8^+^ T cell activation, whereas CD83 loss promotes immunosuppressive T cell phenotypes.

### ALTs promote activation of CD8+ T cells

The former observations uncovered a rare population of ALTs in human glioma that was similarly detected in our pB-IUE model of murine glioma. Because pB-IUE tumors arise *de novo* in an immunocompetent setting, we exploited GFP-labelling of glioma cells and used multispectral imaging flow cytometry to experimentally discern if ALTs can be confirmed *in vivo*. Using this imaging platform, we identified two populations of CD83^+^GFP^+^ cells of interest. The first population featured dendrite-like projections and punctate GFP expression (**Fig. S8A**), which was located both intracellularly and adjacent to CD83 on the cell surface. Past studies employing this imaging modality to examine phagocytosis in live cells^21^ describe this expression pattern as indicative of CD83^+^ immune cells actively phagocytosing or displaying GFP^+^ tumour antigen on their surface. A second population of CD83^+^GFP^+^ cells was characterized by high, diffuse, intracellular GFP expression, a hallmark of targeted cells in the pB-IUE model^22^, and therefore represented SCRAM-identified ALTs (**Fig. 4E; S8B–D**). Subsequent analysis also revealed a rare number of events wherein CD83^−^GFP^−^ cells were bound to GFP^+^ tumour cells at a CD83^+^ interface (**Fig. S8E**), further implicating CD83 as a mediator of tumour–host cell interactions in glioma. These imaging studies confirmed the existence of an endogenous CD83^+^GFP^+^ ALT cell population in the pB-IUE glioma model, similar to the population identified by SCRAM in humans and mice.

Because CD83 is required for antigen presentation in non-tumour immune cells, we examined how changes to tumor cell CD83 expression affect *in vivo* tumour progression. We used CRISPR/Cas9 technology to generate CD83 loss-of-function (CD83^KO^) pB-IUE tumours and separately used murine *Cd83* overexpression to generate CD83 gain-of-function (CD83^OE^) pB-IUE tumours **(Figs. S8F**,**G)**. Kaplan–Meier survival analysis revealed that CD83^KO^ tumours were associated with reduced survival times, whereas CD83^OE^ tumours were associated with extended survival compared with control tumours (**Fig. 4F**). Age-matched CD83^OE^ tumours were smaller than control tumours, whereas CD83^KO^ tumours appeared significantly larger (**Figs. 4G,H**). Cell proliferation assays using Ki67 showed that CD83^OE^ tumours were less proliferative than age-matched CD83^KO^ and control tumours; however, no difference in proliferation was observed between control and CD83^KO^ tumours (**Figs. S8H**,**I**). Considering the worse survival outcomes associated with CD83^KO^ tumours, we analysed the number of nuclei per tumour field of view and found that CD83^KO^ tumours contained nearly 50% more cells than control tumours (**Fig. S8J**), raising the possibility that impaired immune clearance of tumour cells may account for CD83^KO^ phenotypes. These collective observations demonstrate that intra-tumoural CD83 expression modulates disease progression in mouse glioma and suggest that tumour extrinsic mechanisms may contribute to poor survival outcomes when CD83 expression is lost.

CD83 directs T cell responses by enhancing MHC gene expression and enabling antigen presentation by APCs^23^; therefore, we characterized how changes in CD83 expression levels in tumour cells affect T cell responses in glioma. We derived glioma cell lines from pB-IUE tumours and performed co-culture experiments with naïve CD8^+^ T cells (**Fig. 4I**). Following 1 week of co-culture, T cells were harvested for imaging flow cytometry, which revealed that CD83^OE^ co-cultures had the largest CD3^+^CD8^+^ population (**Fig. 4J**). Closer examination uncovered a subset of CD3^−^CD8^+^ cells in CD83^KO^ co-cultures, greater than 90% of which expressed the naïve T cell marker C–C chemokine receptor type 7 (CCR7) and greater than 76% of which co-expressed the activated marker CD25 (**Fig. 4K**). Prior studies show that CD3 loss reduces T cell expansion^24,25^, and CCR7 loss is required for the acquisition of cytotoxic functions in CD8^+^ effector memory T cells^26,27^. Additionally, a subset of CD8^+^CCR7^+^CD25^+^ T cells has been implicated as potent immunosuppressive mediators^28^, suggesting that CD3^−^CD8^+^CCR7^+^ T cells may contribute to subpar T cell responses in CD83^KO^ tumours. Consistent with these data, enzyme-linked immunosorbent assay (ELISA) analyses of co-culture media revealed that both CD83^OE^ and CD83^KO^ co-cultures showed increased production of the T cell–activating cytokines interferon-gamma (IFNγ) and tumour necrosis factor-alpha (TNFα)^29,30^, but only CD83 ^OE^ co-cultures showed increased interleukin (IL)-2, which is required for memory and effector T cell subsets^31^ (**Figs. 4L,M**). Our scRNA-seq analyses of CD83-altered tumours confirmed that CD83^OE^ tumours are enriched for T cell–activating cytokines (**Fig. 4N**) and that T cells from CD83^KO^ tumours displayed increased expression of anergic and regulatory T cell markers. In contrast, T cells from CD83^OE^ tumours showed high expression of *CD44*, a prominent memory and effector T cell activation marker^32^ (**Fig. 4O**). Notably, *in vitro* 5-ethynyl-2′-deoxyuridine (EdU) assays revealed that CD83^OE^ tumour cells were intrinsically more proliferative than control tumour cells and had similar proliferation rates to CD83^KO^ tumour cells (**Figs. 4P,Q**). We confirmed this result through scRNA-seq cell cycle scoring of CD83-altered tumours, which showed increased expression of proliferation markers in CD83^OE^ and CD83^KO^ tumour cells compared with control tumour cells (**Figs. S8K–M**). Combined with our *in vivo* proliferation and survival study results, these findings suggest that T cell–CD83 interactions in CD83^OE^ tumours promote tumour cell clearance and counteract glioma cell proliferation, thus ameliorating poor survival outcomes (**Fig. 4R**). These results demonstrate that modulation of CD83 expression in tumours can direct glioma progression through both cell autonomous proliferation effects and cell non-autonomous responses mediated by T cell–dependent changes to the cytokine milieu. Moreover, these studies emphasize the ability of SCRAM to identify rare, co-occurring tumour cell populations that may contribute to tumour biology.

### Functional neuronal-like tumour cells are unique to *IDH1* mutant glioma

Parallel analyses using SCRAM revealed that most tumour cells from IDH1^mut^ samples mapped to Cluster 4, which displayed a high density of *IDH1* mutations and featured chromosome 1p19q co-deletions in oligodendroglioma patients (**Figs. 5A–C**). We found that 40% of cells (4324 out of 10739 cells) with CNV or GSC annotation had co-occurring cell type annotations of mature neurons, GABAergic neurons and/or synaptic neurons, as well as oligodendrocytes and OPCs (**Fig. 5D**); in contrast, only 18 of 349 tumour cells had co-occurring neuronal signatures in IDH1^WT^ samples. SCRAM identified a similar population of neuronal-like tumour cells (NLTs) in our mouse glioma dataset (**Figs. S9A**,**B**), which prompted us to further investigate these cell types.

**Figure 5.**
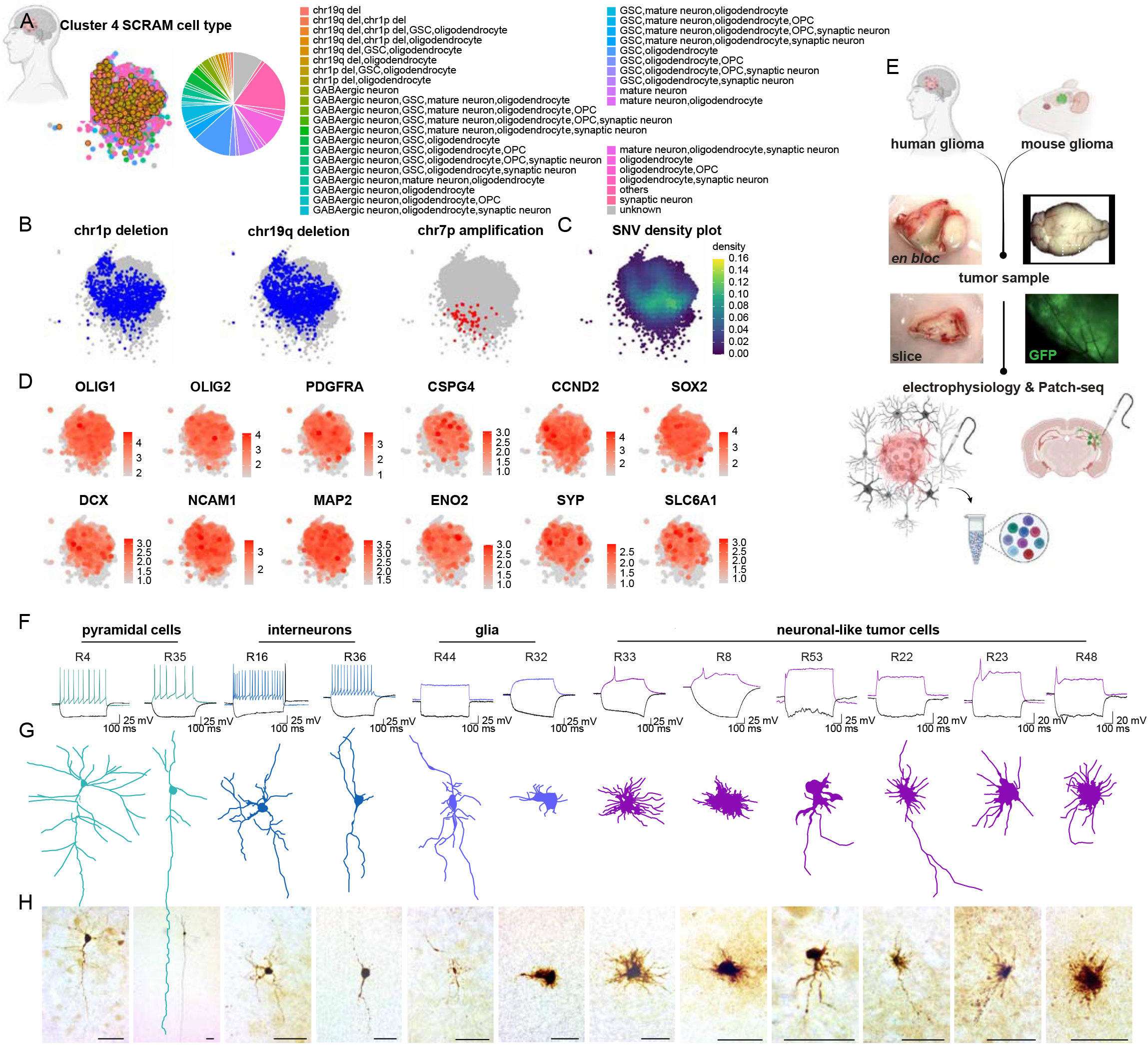
Neuron-like tumour cells are specific to *IDH1*-mutant glioma and functionally similar to neurons. (A) Dimplot and corresponding pie chart showing Single-Cell Rule Association Mining (SCRAM)-identified neuron-like tumour cells (NLTs) for Cluster 4 in human glioma samples. Copy-number variant (CNV)-harbouring cells are outlined in black. (B) Feature plots showing exemplary large-scale CNVs. (C) Density plot showing single-nucleotide variant (SNV) density. (D) Feature plots for select GSC (*CCND2, OLIG2, SOX2*), oligodendrocyte (*OLIG1, OLIG2*), OPC (*PDGFRA, CSPG4*), mature neuron (*MAP2, ENO2*), synaptic neuron *(MAP2, ENO2, SYP*), and GABAergic neuron (*DCX, NCAM1, SLC6A1*) markers. (E) Experimental workflow for whole-cell recordings and whole-cell patch clamp recordings, followed by singe-cell RNA-sequencing (Patch-seq) assays. (F) Exemplary membrane responses from patched *IDH1-*mutant (IDH1^mut^) cells to a 600-ms hyperpolarizing current step (black) and suprathreshold depolarizing current step (coloured). Two voltage traces are shown: the hyperpolarization trace obtained with injected currents (black) and the depolarization trace showing maximal AP firing rate; injected current: −100 pA. (G) Matched traced cell morphologies are shown for recorded cells. (H) Matched images of biocytin-filled cell morphologies for patched cells; scale bar: 50 µm. AP: action potential; GABA: γ-aminobutyric acid; GSC: glioma stem cell; OPC: oligodendrocyte precursor cell.

To validate the existence of NLTs, we performed whole-cell patch clamp recordings, followed by scRNA-seq (Patch-seq), on brain slices resected from the tumour cores of two IDH1^mut^ glioma specimens (**Fig. 5E**). We recorded from 53 total cells; 31 were used to recover high-quality RNA for scRNA-seq, and 20 were preserved with biocytin for morphological analysis (**Figs. 5F–H; Figs. S10A–H**). Of these cells, 27 displayed electrophysiological and morphological properties consistent with either glia or neurons in the non-tumour brain^33,34^, whereas 26 cells displayed select neuronal electrophysiological properties but were morphologically consistent with glia, and therefore were presumed to be NLTs. These presumptive NLTs had higher input resistances than either typical neurons or glia (**Fig. 6A**) and had the capacity to fire single, small action potentials (APs; **Figs. 6B,C**), signifying that the acquired transcriptional signatures of these cells correlated with neuronal functional characteristics. Prior work describes similar electrophysiological properties for early-stage, post-mitotic, newborn neurons, implicating hyperexcitability, high input resistance, and the firing of single APs as defining features of this developmental stage^35^. Newborn neurons also feature high GABA receptor levels^36,37^, which are also expressed in NLTs, and SCRAM identified several cells with annotations for both GABAergic neurons and IDH1^mut^ tumour cells. Our Patch-seq results revealed the presence of the canonical *IDH1R132H* mutation in five of 15 NLTs, confirming that these cells were tumour (**Fig. 6D; Figs. S11A**). Subsequent cell type enrichment analysis of Patch-seq data using the EnrichR tool^6,38^ demonstrated that NLTs are transcriptionally similar to glia, showing enrichment for embryonic, astrocytic, and OPC gene sets (**Figs. 6E,F**). Using the SCRAM *de novo* marker tool, we identified *SOX11* as an NLT marker in both Patch-seq and scRNA-seq datasets (**Figs. S11B**,**C**). When we attempted to record from an IDH1^WT^ glioblastoma core sample, very few cells were patchable due to gliosis and necrosis; however, all three cells recorded were electrophysiologically consistent with glia (**Fig. S10H**). Using *in vivo* modelling, whole-cell recordings were obtained for GFP^+^ tumour cells from control pB-IUE model mice, revealing an analogous population of NLTs (**Figs. S11D**,**E**). Taken together, these human and mouse studies confirm the existence of NLTs featuring transcriptional and electrophysiological properties similar to but distinct from neurons as a defining feature of human IDH1^mut^ glioma.

**Figure 6.**
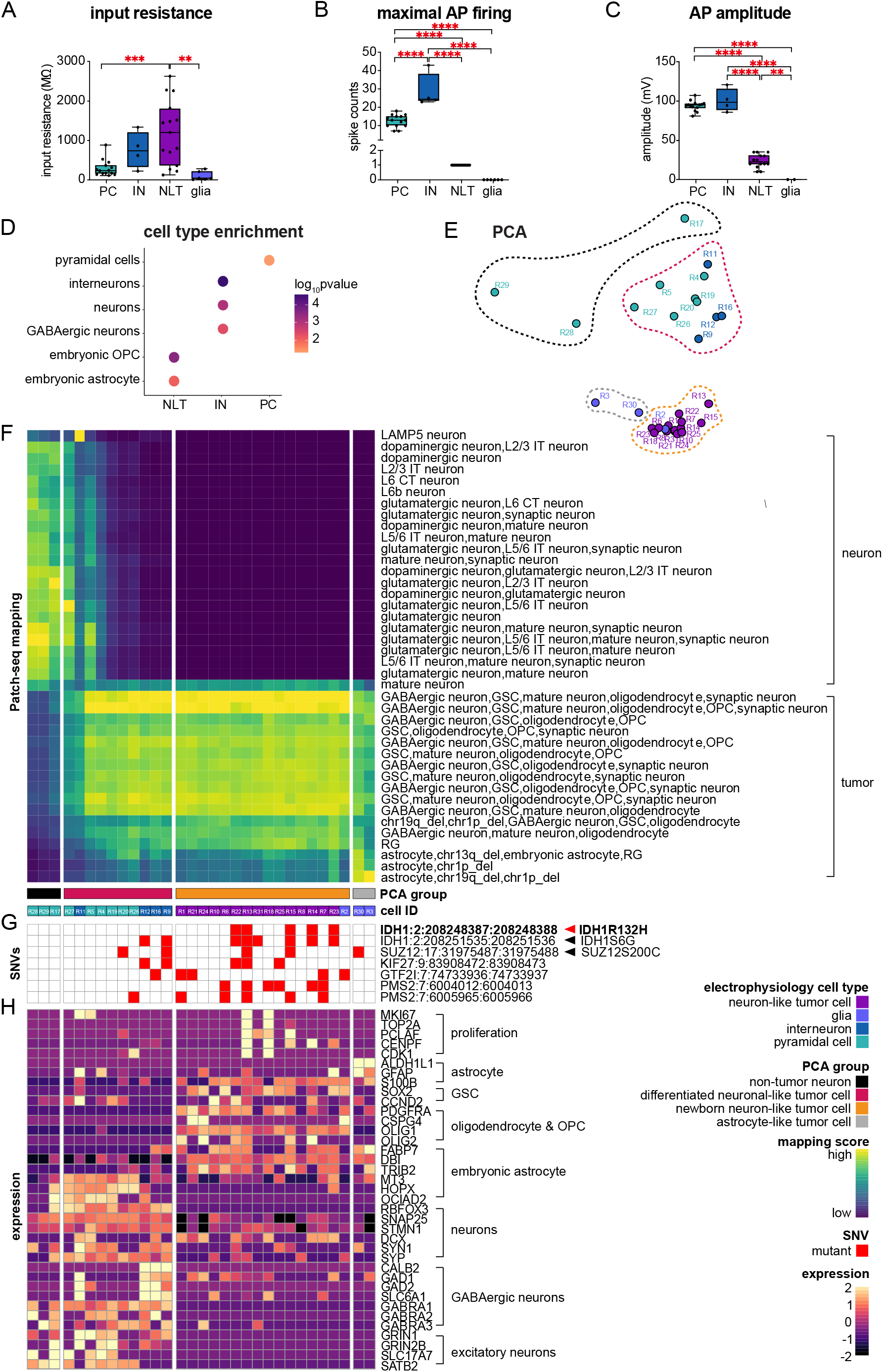
Neuron-like tumour cells are a novel cell type with unique electrophysiological, transcriptomic and genomic profiles. (A) Bar graph showing neuron-like tumour cells (NLTs) have higher input resistance compared with non-tumour neurons; *p*-values for pairwise comparisons are noted in the figure. PC: pyramidal cell; IN: interneuron. (B) Bar graph showing NLTs fire fewer spikes compared to neurons; *p*-values for pairwise comparisons are noted in the figure. (C) Bar graph showing NLTs have smaller action potential (AP) amplitudes compared to neurons; *p*-values for pairwise comparisons are noted in the figure. (D) EnrichR analysis of 31 cells analysed by whole-cell patch clamp recordings, followed by single-cell RNA-sequencing (Patch-seq), confirms NLTs are enriched for gene sets corresponding to oligodendrocyte precursor cells (OPCs) and embryonic astrocytes; *p*<0.05. (E) Principal component analysis (PCA) plot of 31 Patch-seq cells shows clustering of cells based on electrophysiological properties. Newborn NLTs are transcriptionally similar to each other and glia, whereas differentiated NLTs resemble non-tumour pyramidal cells. Cells are coloured according to electrophysiological properties; dotted lines show PCA groups. (F) Mapping Patch-seq cells to Single-Cell Rule Association Mining (SCRAM)-annotated single-cell RNA-sequencing human glioma dataset and (G) single-nucleotide variant (SNV) analysis shows newborn NLTs map strongly to NLT annotations, whereas differentiated NLTs map to both non-tumour neurons and NLTs. Astrocyte-like tumour cells show reduced NLT profiles and map most strongly to astrocyte-like, chromosome 1p19q co-deleted tumour cells from *IDH1-*mutant (IDH1^mut^) oligodendroglioma. (G) The top seven most frequently occurring SNVs are shown for each cell. (H) Heatmap showing expression of representative genes from neural, glioma, and proliferating cell types. Proliferation markers are only detected in newborn and differentiated NLTs. GSC: glioma stem cell; RG: radial glia.

We mapped the 31 Patch-seq cells to our human glioma scRNA-seq dataset and found that all 15 NLTs examined by Patch-seq mapped to GABAergic neurons with co-occurring GSC, OPC, or oligodendrocyte annotations (**Figs. 6F,G**). Although only five Patch-seq NLTs presented with the canonical *IDH1R132H* mutation, 12 NLTs presented with *SUZ12, KIF27, GTF2I, PMS2*, or non-canonical *IDH1* mutations (**Figs. S11F**,**G**), all of which were previously identified in both IDH1^WT^ and IDH1^mut^ patients in our scRNA-seq dataset (**Fig. S3H**). Using Patch-seq mapping, together with principal component analysis (PCA) and SNV analysis, we divided the 31 Patch-seq cells into subclasses based on electrophysiology profiles, PCA clustering, mapping to our SCRAM-annotated scRNA-seq dataset, SNV incidence, and cell type marker expression (**Figs. 6E–J; Fig. S12**). Employing this strategy, we uncovered 10 cells with electrophysiological properties similar to either pyramidal cells or interneurons that mapped strongly to NLT-annotated cells, five of which featured similar mutational profiles to NLTs. Unlike early-stage newborn NLTs, which are electrophysiologically distinct from differentiated interneurons and pyramidal cells, these mutation-bearing pyramidal cell– and interneuron-like cells, which we refer to as differentiated NLTs, were not electrophysiologically distinct from non-tumour neurons; however, they trended towards input resistances and AP amplitudes between those of non-tumour neurons and newborn NLTs, and had high maximal AP firing rates (**Figs. S13A–C**). In addition, differentiated NLTs were morphologically similar to non-tumour neurons but mapped to both non-tumour neuronal cell types and NLTs in our scRNA-seq dataset, whereas newborn NLTs did not show strong neuronal transcriptional profiles. Examination of Patch-seq transcriptional profiles revealed proliferation markers were only expressed by newborn and differentiated NLTs, suggesting that NLTs represent the major proliferative constituents in these tumours (**Fig. 6J**). We found that embryonic astrocytic genes were enriched in differentiated NLTs, which also feature high expression of RNA polymerase III, the overexpression of which has been linked to DNA damage and cancer^39,40^ (**Fig. 6J; Figs. S13D**,**E**). These observations suggest that more than one NLT cell type exists and that these NLT cell types collectively form a neurodevelopmental spectrum paralleling the processes of neuronal differentiation. Whether differentiated NLTs represent astrocyte-derived tumour cells that have acquired the transcriptional and electrophysiological properties of differentiated neuronal subtypes or are indicative of non-tumour neurons undergoing transcriptomic remodelling and transformation remains an important question, the answer to which could shed light on the origins of glioma cells and should be further explored in future studies.

## Discussion

### A computational framework for defining genomic and transcriptional states

By 2040 the number of new cancer cases per year will be 29.5 million worldwide, more than half of which will end in cancer-related deaths^41^. While advances in single-cell sequencing technologies have given scientists the opportunity to examine individual cells in unprecedented detail, the power of these data has yet to be harnessed by computational platforms capable of resolving these profiles on a cell-by-cell basis. To address this unmet need, we developed SCRAM, which we report here as the first computational algorithm of its kind capable of defining integrated co-occurring, genomic and transcriptomic features from single cell datasets. In this report, we employed SCRAM using a curated brain tumour-specific cell type marker list to elucidate two previously unreported glioma cell types; however, our pipeline can be easily expanded for use in any cancer, non-oncologic, or developmental context and thus represents a new computational platform that can be used to study human and mouse biology.

### ALTs enable antitumor T cell responses

Owing in part to the existence of a highly immunosuppressive tumour microenvironment, immunotherapy trials for glioma have been largely unsuccessful, with little to no overall survival benefits^42,43^. Prior tumour immunology research has characterized the molecular mechanisms mediating this immunosuppressive environment in glioma, implicating anergic and regulatory T cells as important mediators^44,45^. Researchers have shown that tumour cells present endogenous antigens via MHC, and speculated that MHC loss in cancer impairs naïve T cell priming and subsequent antitumour cytotoxic T cell responses^46–50^. Considered alongside prior work confirming CD83 as a regulator of MHC expression^51,52^, our data implicate CD83 as a mediator of antigen presentation by glioma cells and suggest that CD83 loss promotes immunosuppressive T cell responses by impairing glioma cell antigen presentation. Our imaging flow cytometry experiments confirmed that GFP^+^ tumour cells are actively surveilled by the surrounding immune environment, implicating downstream immunosuppressive responses as likely culprits for poor survival outcomes. Whether changes to the proportions of activated CD8^+^ T cells in CD83-altered tumours correlate with their abilities to recognize and destroy glioma cells remains unanswered and should be the focus of future inquiries.

### NLTs are a hallmark of IDH1^mut^ glioma

Over the past decade, the emerging field of cancer neuroscience has elucidated how glioma cells interact with surrounding neural networks to direct disease progression^53,54^. Prior studies have reported that human glioma cells receive synaptic inputs from the surrounding neuronal circuitry^55–57^. Our studies build upon past reports by suggesting that SCRAM-identified NLTs may communicate with peri-tumoural neurons by directing outgoing electrochemical information in the form of APs. These data suggest that dysregulated electrophysiological activity in the glioma-bearing brain may originate from tumour cells able to initiate and propagate outgoing neurochemical signals. Importantly, NLTs were the only cells expressing proliferation markers in our Patch-seq dataset, suggesting that NLTs are important contributors to both the neurophysiological and proliferative characteristics of glioma. Clinically, IDH1^mut^ glioma frequently presents with seizures^58^, raising the possibility that NLTs may contribute to both hyperexcitable and proliferative phenotypes in these tumors.

Our studies also uncovered a SNV-bearing population of neurons featuring transcriptomic profiles consistent with both non-tumour neurons and tumour cells, which we termed differentiated NLTs. The existence of these cells brings into focus the longstanding and controversial topic of tumour cell origins. Prior work has theorized that GSCs arise from OPCs^59–61^, which is supported by our observations that newborn NLTs are enriched for OPC and GSC signatures. However, by layering our SNV analysis onto both transcriptional and electrophysiological profiling, we found that differentiated NLTs have a lower mutational burden than other tumour cells. These data imply that differentiated NLTs may not derive from the tumour but may instead represent a population of peri-tumoural neurons undergoing malignant transformation. Future investigations remain necessary to confirm the origins of differentiated NLTs, which will improve the current understanding of how tumor cells arise in malignant glioma.

## Acknowledgements

This work was supported by grants from the NIH (R01NS071153 to B.D., R01NS124093 to B.D. and G.R., R01CA223388 to B.D, R01NS094615 to G.R.). R.N.C. is supported by grants from the NIH (5T32HL92332-15, F31CA265156, and F99CA274700). The Baylor College of Medicine Single Cell Genomics Core is supported by NIH Shared Instrument Grants (S10OD023469, S10OD025240), P30EY002520, and CPRIT grant RP200504. Schematics were created using Biorender.com. The Cytometry and Cell Sorting Core at Baylor College of Medicine is supported by funding from the CPRIT Core Facility Support Award (CPRIT-RP180672), the NIH (CA125123 and RR024574), and the assistance of Joel M. Sederstrom. BioSorter is supported by NIH grant S10 OD025251.

## Data availability

The Patch-seq and scRNA-seq datasets generated during this study will be made available through the NCBI Gene Expression Omnibus (GEO) website. All other study data are included in the article and/or supporting information.

## Code availability

The source code of SCRAM will be available upon publication.

## Author contributions

R.N.C. and A.S.H. are responsible for conception of this project, the study and pipeline design, and interpretation of the results. R.N.C. performed all pB-IUE surgeries, *in vitro* and FACS experiments, immunostaining, and statistical analyses and prepared all scRNA-seq samples from humans and mice with assistance from M.F.M., and A.S.H. wrote the code for the SCRAM pipeline. Q.M. and J.J. performed all human electrophysiology and Patch-seq experiments. J.M. performed the two-photon and widefield imaging studies in mice and subsequent analyses. I.A. performed all mouse electrophysiology and respective analyses. R.N.C., A.O.H., and A.S.H. performed bioinformatics analyses on human and mouse scRNA-seq datasets. A.C. maintained all mouse colonies and assisted with tissue harvesting and fixation. B.L., Z.L., D.C, E.L.F., Y.K., P.H., and A.R. provided experimental support for *in vitro* cell culture, immunohistochemistry, immunofluorescence, immunocytochemistry, and surveyor experiments. C.M. performed histopathological diagnoses of mouse tumours. A.J. and G.R. identified and obtained consent from patients for the study. R.N.C. and A.S.H prepared the manuscript. J.N., X.J., B.D., and G.R. contributed to the manuscript with feedback from all authors.

## Contact for reagent and resource sharing

Dr. Akdes Serin Harmanci is the lead contact for reagent and resource sharing. All published reagents will be shared on an unrestricted basis; reagent requests should be directed to the corresponding author.

